# Phylogenetic, sequence and structural analysis of Insulin superfamily proteins reveals an indelible link between evolution and structure-function relationship

**DOI:** 10.1101/769497

**Authors:** Shrilakshmi S., Shankar V. Kundapura, Debayan Dey, Ananda Kulal, Udupi A. Ramagopal

## Abstract

The insulin superfamily proteins (ISPs), in particular, insulin, IGFs and relaxins are key modulators of animal physiology. They are known to have evolved from the same ancestral gene and have diverged into proteins with varied sequences and distinct functions, but maintain a similar structural architecture stabilized by highly conserved disulphide bridges. A recent surge of sequence data and the structures of these proteins prompted a need for a comprehensive analysis which connects the evolution of these sequences in the light of available functional and structural information and their interaction with cognate receptors. This study reveals a) unusually high sequence conservation of IGFs (>90%), which has never been reported before. In fact, it was interesting to observe that the functional domains (excluding signal peptide) of human, horse, pig and Ord’s kangaroo rat are 100% identical. (b) an updated definition of the signature motif of the relaxin family (c) a non-canonical C-peptide cleavage site in a few killifish insulin sequences and so on. We also provide a structure-based rationale for such conservation by introducing a concept called binding partners imposed evolutionary constraints. Furthermore, the high conservation of IGFs appears to represent a classic case of resistance to sequence diversity exerted by physiologically important interactions with multiple partners. Furthermore, we propose a probable mechanism for C-peptide cleavage in killifish insulin sequences.

## Introduction

Insulin superfamily proteins (ISPs) are involved in critical functions and few of which are extensively studied considering their therapeutic applications. The insulin superfamily includes many homologous proteins such as Insulin (1), IGF-1 (insulin-like growth factor-1) (2), IGF-2 (3), Relaxin (Relaxin 1 2 &3) (4), Insulin-like peptides (INSL3, INSL4, INSL5, INSL6, ILP-1 and ILP-2)(5,6), Bombyxin (7), Locust insulin related peptides (8), Molluscan insulin-related peptides (9) and Caenorhabditis elegans insulin-like peptides (10), which are found in most of the animal phyla. ISPs share similar structural architecture with a distinct insulin fold formed by three conserved disulphide linkages. All the insulin superfamily proteins and peptides are proposed to be evolved from a single ancestral gene after undergoing conducive multiple independent gene duplications (11).

Insulin is the most studied member of this family. Considering its importance in glucose metabolism and diabetes, it is probably most explored or a protein experimentally “beaten to death”. Since, ISPs have evolved from a single ancestral gene, we asked the question whether the collective analysis of these members with respect to their sequence divergence/similarity, structure, function and their known interaction with cognate partner(s) reveal any novel insights? With the above question in mind, we have analyzed insulin, insulin-like growth factors (IGFs), bombyxin and relaxin. Among the four members of ISPs considered here, three of these proteins are found in humans (insulin, IGFs and Relaxins) whereas, Bombyxin is an insect protein. Since this study was aimed at understanding the evolution of ISP members in relation to their structure and function, we chose the above mentioned four families of ISPs as these members are relatively well explored, either biochemically and/or structural information on these molecules individually or in complex with their receptors are available. Although, in the last few decades many structure, evolution and functional studies (12–15) have been performed, to our knowledge, there is no study that involves sequence, structure, function and phylogenetic perspective encompassing all the four major families of insulin superfamily. With the surge of sequence information in the post genomics era and recently available structures of several of these ISP members (16–20) in few cases their complexes with receptors, we were curious to know, whether any novel insights could be derived from the cumulative analysis of these evolutionarily related but functionally diverse molecules having similar structural architecture.

As mentioned above, insulin is the well-known member of this family. In brief, this miniature protein is secreted by the pancreatic β cells. The main function of insulin is to maintain optimal blood glucose levels by facilitating glucose uptake by cells, which is mediated by interaction of insulin with its receptor (IR). It also plays a vital role in gene regulation and metabolism of proteins, carbohydrates and lipids (21).

Insulin-like growth factors (IGF-1 and IGF-2) have important roles in regulating the growth and metabolism of all cells and are primarily produced by the liver as a endocrine hormone (2). Unlike insulin, IGFs retain their C-peptide (also called C-domain) and are single polypeptide chains which maintains the insulin fold. IGF-1 and IGF-2 have similar functional roles except that they are expressed at different stages of physiological development. IGF-1 is expressed mainly after birth whereas IGF-2 is expressed during early embryonic and fetal development in most of the somatic cells (22).

Relaxins are a family of proteins in insulin superfamily having many paralogs among them like relaxin-1, relaxin-2, relaxin-3 and so on (23). Relaxin-2 is known to mediate the hemodynamic changes that occur during pregnancy and is pivotal in mammals which relaxes pelvic ligaments, hence the name relaxin (24). Besides its role in pregnancy, Relaxin-2 also plays a key role in vasodilation, antifibrinolysis, wound healing and is recently found to be involved in cardiovascular regulation against acute heart failure (25–27). Relaxin-1 is considered to be a homolog of relaxin-2 in primates (5). In humans, relaxin-2 is a major circulating hormone (28) and is expressed in variety of tissues whereas, expression of relaxin-1 is restricted to decidua, placenta and prostrate (5). Relaxin-3 is a neuropeptide hormone found in the brain which is said to have a role in learning, stress, alertness, memory, locomotion and appetite regulation (16, 26). It is also claimed to act as a neurotransmitter (29). Relaxin-3 is widely accepted to be the ancestral gene in relaxin family (5).

Bombyxin is largely found in the lepidopteron class of insects. Bombyxin, previously known as prothoracicotropic hormone (PTTH) (7) stimulates the prothoracic glands to synthesize and release ecdysone, which is a key hormone that regulates insect moulting and metamorphosis (30). It also has a role in trehalose sugar metabolism (31). Like relaxins, bombyxins are known to have many paralogs, although the functions of most paralogs have not been determined experimentally.

Considering the fact, that ISPs have evolved from a single ancestral gene, it is possible that the evolutionary conservation and variations of residues among ISPs, could provide critical clues for rational modification of insulin. The study also attempts to correlate the mode of interaction of ISPs to their respective receptors, the determinants responsible for oligomerization observed in few members of ISPs and the consequent evolutionary restraints. During the course of this analysis we came across few interesting aspects such as (a) unusually high sequence conservation in IGFs for which we attempt to provide a rationale. (b) evolutionarily diverged insulin killifish sequences, with a distinct non-canonical C peptide cleavage site (c) an unnoticed conservation in the receptor binding motif of relaxins, particularly in relaxin-3 and so on. Finally, we have borrowed information from the ISPs to design few plausible insulin analogues, computationally analyzed their structural stability and their affinity to insulin receptor.

## Results and discussion

Insulin is synthesized as a single polypeptide consisting of (a) a N-terminal stretch called B-chain consisting of around 30 residues (b) a post-translationally cleavable stretch consisting around 30 residues called C-peptide and (c) a C-terminal stretch consisting of around 21 residues called A chain (Figure S1) (32). This naming convention is common to all the members of this family, even for the members that do not require post-translational cleavage of C-peptide. We maintain similar convention all through the manuscript. Most of our analysis considers variation and similarities in the B and A chain unless otherwise mentioned.

Insulin superfamily proteins (ISPs) have varying sequence identity amongst themselves (Table S1), yet have a similar structural architecture consisting of three uniquely arranged helices forming the “insulin fold” (12), stabilized by completely conserved three disulfide bridges (Figure S2). For example, in humans, insulin shares a sequence identity of ~30% with IGF-1 and its sequence identity with relaxin-2 is as low as 14%. All the three members belong to functionally divergent family (Table S1), suggesting, in ISPs a sequence identity of below 30% probably represent two sequences of divergent function. Similarly, when the two IGF paralogs having similar function (both are growth hormones) were compared they had a sequence identity 63% between them. Further, within the relaxin family, Relaxin-1 shares a sequence identity of ~80% with its functionally equivalent member Relaxin-2 (both play a role in pregnancy and vasodilation), whereas, its sequence identity with functionally distinct paralog, a neuropeptide, relaxin-3 is as low as 23%. The above observations suggest that the sequence similarities/differences, although indicative of their function, are not useful to distinguish ISP members into different families.

We have used minimum evolution as the phylogenetic method to understand the relationship between the proteins of the insulin superfamily. When the insulin superfamily phylogenetic tree (Figure 1) was generated, it clearly diverged into four families (Insulin, IGFs, bombyxin and relaxin). The sequences obtained from the clades of these families were considered to recognize position specific conservation and variation within a given family and amongst the families.

**Figure 1:**
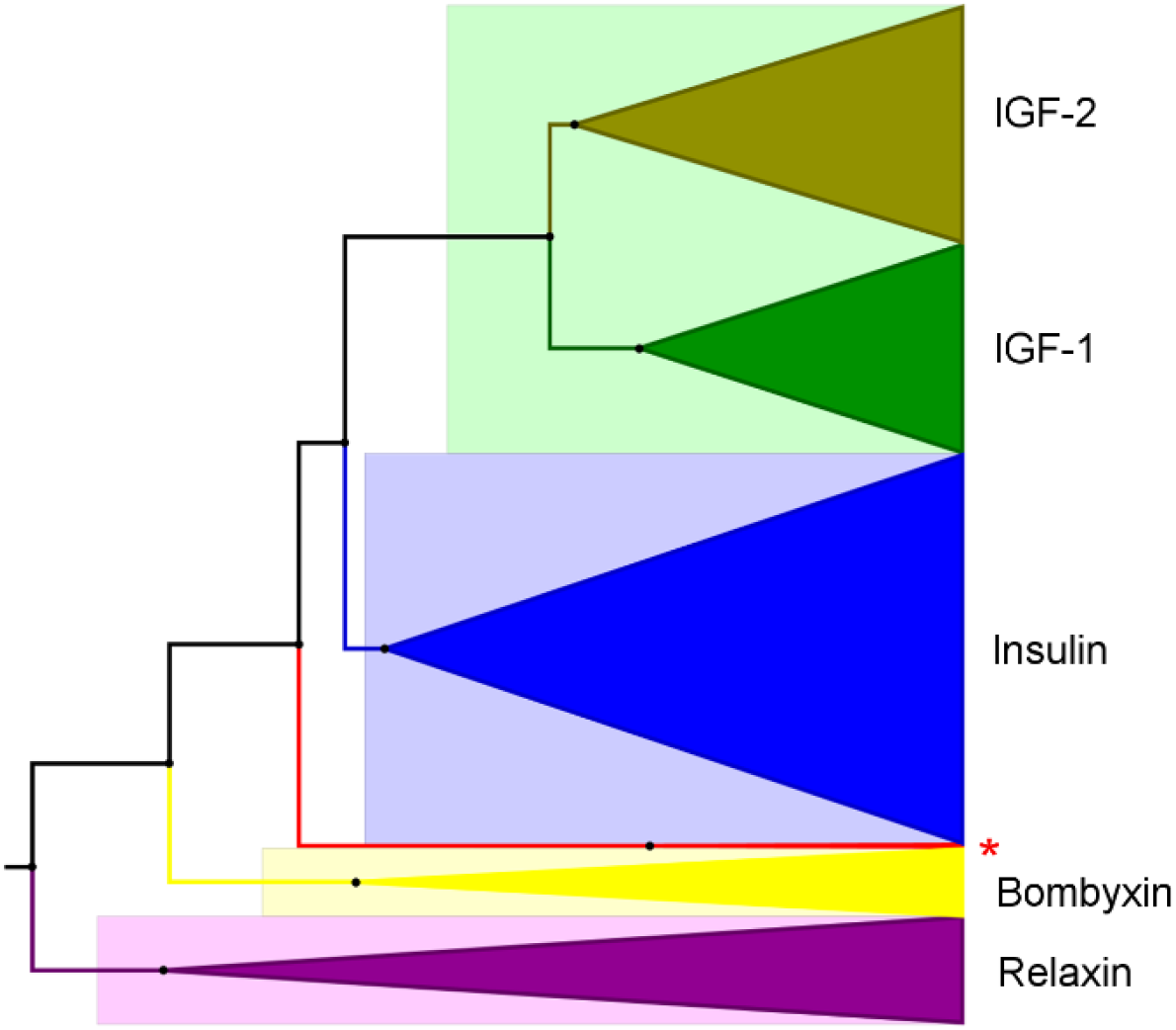
Phylogenetic tree of ISPs. constructed using minimum evolution and visualized by graphical tool FigTree. Green clade represents IGF-1, dull green clade represents IGF-2, blue clade represents insulin family, yellow clade represents bombyxin family and purple clade represents relaxin family. The star represents a separate clade of divergent killifish insulin sequences.

The phylogenetic tree (Figure 1) suggests that there was an early divergence of relaxin family from other ISPs. The next family to diverge was bombyxin which consists of sequences largely from *Bombyx mori*. A small clade diverges out before insulin and IGFs which will be discussed later in section on C-peptide cleavage site of killifish insulins. A common ancestor of insulin and IGFs is observed after which insulin and IGFs diverge out together. Further, two clades diverge from IGFs in to IGF-1 and IGF-2. The phylogenetic analysis indicates that the insulin superfamily proteins are closely related to each other with an exception of relaxin which is distant and diverges out early. It is also noteworthy to indicate that, each family has undergone gene duplication events amongst themselves and thereby have different paralogs in the same family.

The sequences clustered based on phylogenetic analysis were analyzed for conservation within each family and among all the ISP members. The sequence logo generated for each family and the common sequence logo for all the ISP members called “master sequence” is represented in the figure 2. It is clear that B8Gly and the cysteines involved in the formation of canonical disulphide linkages are absolutely conserved throughout the family (Figure 2, black asterisks). In insulin, the B chain has a higher conservation than A chain, which is consistent with the previous report (14). Insulin is the only protein of this family which is stored as a hexamer in its stable form. Hexamerization is stabilized by few hydrophobic interactions between the insulin dimers together with zinc mediated trimerization of insulin dimers, where B10His of every monomer coordinates with the zinc ion to form a hexamer (32). Analysis of the structures of insulin and its complex with its receptor reveal that the highly conserved residues, other than cysteines (see Figure 2) participate in either oligomerization and/or are involved in receptor binding.

**Figure 2:**
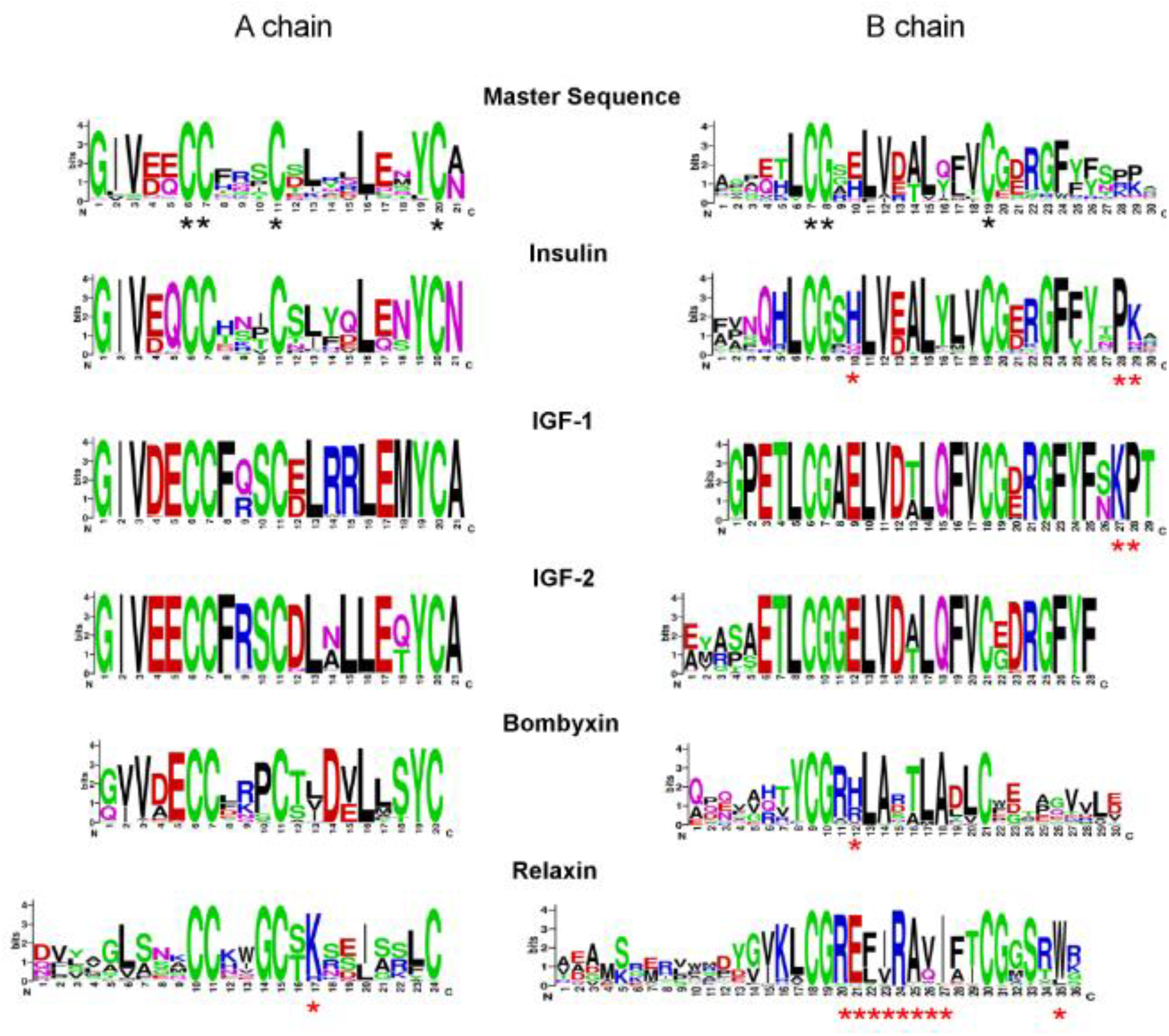
Sequence logos of A and B chains of insulin superfamily. Master alignment sequence logo represents the whole data set whereas; the individual sequence logo of proteins represents their respective cluster (family). The conservation of the residues corresponds to the height of the letter and the sequence logo was generated using the program WebLogo. For convenience, sequence logos of individual protein families are stacked upon each other in such a manner that the conserved canonical cysteines are in the same position. The asterisks in black below the master sequence represents the completely conserved residues in the ISPs. The asterisks in red represent the residues will be discussed in the following sections.

IGFs (IGF-1 and IGF-2) are monomers and they bind to their respective cognate receptors (IGF-1R and IGF-2R) and have low affinity cross reactivity with insulin receptors (IR). From figure 2, it is clear that IGF-1 and IGF-2, despite being monomers, have the most number of completely conserved residues in both A and B chains among ISP members (except for the N-terminal part of IGF-2). This unusual conservation of IGFs, to our knowledge has not been noticed before. The rationale for such high sequence conservation in IGFs will be discussed later.

Bombyxin shares around 30% sequence identity with human insulin (Table S1). While the insulin B chain is relatively more conserved than the A chain, our analysis suggests that the bombyxin A chain is more conserved compared to its B chain (Figure 2). Concurrent with this observation, it has been shown that the modification of the A chain reduces the affinity to the bombyxin receptor (33). Unfortunately, the experimental evidence on bombyxin receptor and its interactions are elusive. A report on a probable receptor suggests that there is a 300kDa protein present on the ovarian cells of lepidopteran species (34). It was proposed that bombyxin doesn’t form a hexamer (35, 36) though it contains the B10His (64%), which in insulin plays a vital role in zinc mediated hexamerization.

Relaxins are the first family to diverge according to the phylogenetic tree and have the least sequence conservation when compared to the other families (Figure 2). A plausible reason could be the presence of functionally variant paralogs like relaxin-1, 2 and 3 as mentioned earlier. Although, functionally divergent, relaxin sequences have known to possess a distinct receptor binding motif (RXXXRXXI/V), which is proposed to be important for its interaction with G-protein coupled receptors (GPCRs) (37). Our analysis indicates that this binding motif needs to be redefined with regards to sequence conservation.

### Distinct insulin sequence of Killifish reveals a non-canonical C-peptide cleavage site

As observed in the phylogenetic tree (Figure 1), a small clade diverges out from the ancestral node of insulin and IGFs. This clade consisting of only two insulin sequences belong to the killifish species and share a sequence identity of 39% with human insulin. These sequences belong to Turquoise killifish (Uniprot ID: A0A1A8ACH4) and Beira killifish (Uniprot ID: A0A1A8ISB4). Further, most of the residues involved in oligomerization and receptor binding interactions are maintained when compared to human insulin (Table S2 and S3). Moreover, when the insulin receptor sequences of these killifishes were compared with human insulin receptor sequence, it was observed that they had a sequence identity of around 60%. All the above observations indicate that the insulin from these killifish species might oligomerize and interact with their cognate receptor in a similar manner to human insulin.

The detailed observation of these diverged killifish sequences revealed an interesting anomaly at one of their post-translational C peptide cleavage sites. This cleavage is essential for processing of pro-insulin to its active form by the action of proconvertases. Canonically, the C-peptide cleavage site at the C-terminal end of B chain (cleaved by proconvertase 1/3) and N-terminal end of A chain (cleaved by proconvertase 2), is occupied by two consecutive dibasic residues, consisting of arginine or lysine (figure 3, black asterisk-B chain C-terminal, red asterisk-A chain N terminal). This is classified as type 2 precursor cleavage motif identified by proconvertases (38), which is marked by ((Arg/Lys)-(Arg/Lys)) motif. Interestingly, this dibasic cleavage site at C-terminal end of B chain is replaced by di-acidic aspartate residues (-Asp-Asp-) in the above killifish sequences (Figure 3).

**Figure 3:**
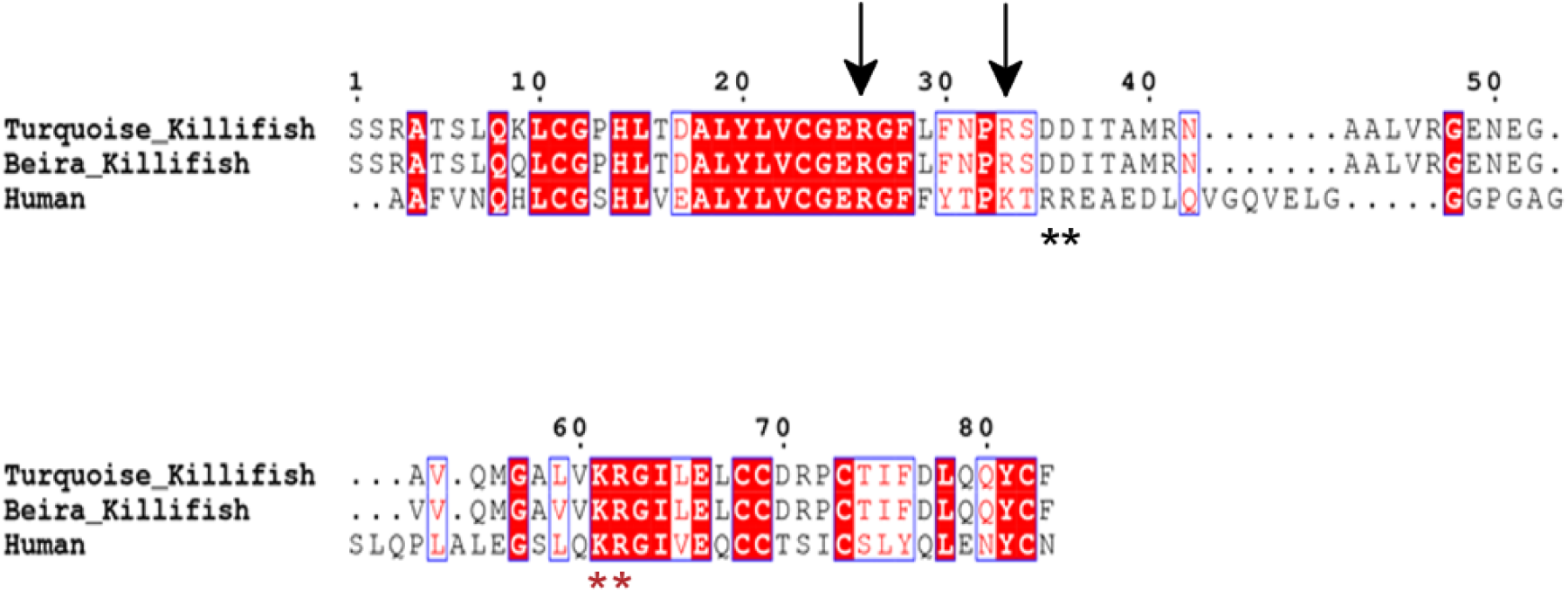
Multiple sequence alignment of insulin sequences, Human insulin (Uniprot ID: P01308), Turquoise killifish insulin (Uniprot ID: A0A1A8ACH4) and Beira killifish insulin (Uniprot ID: A0A1A8ISB4). The two black asterisks represent the change in cleavage site of C peptide at the C-terminal end of B chain. The two red asterisks represent the N-terminal cleavage site of the A chain which are conserved. The black arrows indicate the plausible type 3 cleavage motif found in killifish insulin sequence.

Replacement of two dibasic residues by consecutive residues of identical nature, -Asp-Asp-, requires a change of bases in adjacent codons that lead to two identical residues. Further, the change of codon from Arg to Asp requires a mutation of at least two bases in each codon. All these observations lead to prudent questions about possible C-peptide cleavage or lack thereof. However, upon closer observation, a plausible non-canonical cleavage motif for C-peptide in these killifish insulin was identified. This C-peptide cleavage site seems to adopt a type 3 precursor cleavage motif marked by R/K-X_n_-R motif, where X is any other amino acid and “n” takes integer values of 0, 2, 4, 6 (in killifish sequences n=6, Figure 3) (38). The type 3 precursor cleavage motif is seen in pro-Dynorphin, pro-ANF, and E-peptide cleavage site of pro IGFs, but was never reported in insulin sequences. This suggests that proconvertase 1/3 which probably cleaves the C-peptide at the C-terminal of B chain, does so, by recognizing a different cleavage motif. It is interesting to note that these two killifishes inhabit landlocked fresh water ephemeral pools present mainly in Zimbabwe and Mozambique, and hence the observed non-canonical cleavage site might be the result of location specific evolutionary pressure. This difference in the C-peptide cleavage site of these killifishes might be attributed to their ability to evolve rapidly in response to environmental changes (39–41). In contrast, insulin sequence from a geographically less restricted killifish, Mummichog, found in marine sources of the coast of USA and Canada contains the canonical type 2 cleavage site. It should be noted that the rare type 3 precursor cleavage motif is observed in few other chordates (Figure S3).

### Binding partner(s) imposed evolutionary constraint in IGFs

Another interesting observation to emerge out of this study is the unusually high conservation of IGF family members (IGF-1 and IGF-2), previously unreported. In fact, the functional domain of IGF-1 sequence (excluding signal peptide) of human, horse, pig and Ord’s kangaroo rat are 100% identical.

Unlike IGFs, insulin is stored in its stable hexameric form. Thus, the residues which are important for the stabilization of insulin fold, involved in oligomerization (Table S2), and those that interact with the cognate receptor (Table S3) are expected to be conserved during the evolution. However, IGFs are known to be monomers and only obvious evolutionary constraint is on those residues that are involved in interaction with their cognate receptors (IGF-1R and IGF-2R), suggesting IGFs should have less restraints on residue conservation. Surprisingly, among the ISPs, IGFs have the highest number of conserved residues in the A and B chains. In IGF-1, among the 50 residues of A and B chains, around 45 residues have conservation above 98% (Figure 2). Similarly, in the case of IGF-2 among the 49 residues, 37 of them have conservation above 98% (Figure 2).

It is known that the action of IGFs is regulated by IGF-binding proteins (IGFBPs) (42). IGFBPs bind to IGFs and increase the half-life of IGFs in the circulation (43) and regulate receptor binding, thus regulate their signaling (42). There are six IGFBPs (IGFBP-1 to IGFBP-6), they bind to IGFs with equal or greater affinity than the IGF-1R (44). Further, among ISPs, IGFs are the only two members whose interaction with their receptors is known to be controlled by regulatory proteins. The interaction of IGF-1 with its receptors and IGFBPs is depicted with a simple interactome (Figure S4). Based on these observations, we speculate that the observed high conservation of IGFs is probably due to the constraints imposed by its cognate receptors and several of these IGFBPs.

Through biophysical and mutational studies, it has been shown that the N and C-terminal domains of the IGFBPs are involved in binding to IGFs (45). Later, it was confirmed by the crystal structure of the complex of IGF-1 together with N and C-terminal domains of IGFBP-4 (46). Similarly, crystal structure of the complex of IGF-1 in combination with C-terminal domain of IGFBP-1 and N-terminal domain of IGFBP-4 also confirmed the similar mode of binding of IGF-1 with the IGFBPs (PDB entry: 2DSR, 2DSQ) (17). These structures revealed that the IGF-1 binds in a deep cleft formed between the N and C-terminal domains of the IGFBPs (Figure 4A and 4B). Further, buried surface area of IGF-1 upon complex formation (only A and B chain) with IGFBPs, was found to be around 60%. Assuming that the mode of binding of all the IGFBPs (IGFBP1-6) with IGF-1 is similar, even a slight variation in interaction of these IGFBPs with IGF-1 requires burial of additional surface area on IGF-1. Similarly, with additional solvent exposed area of IGF-1 involved in IGF-1R interaction, in particular α-CT region (Figure 4C), there is little room for surface exposed residues to change during the evolution. The above analysis suggests that the N and C-terminal domains of IGFBPs, almost engulf this miniature protein upon complex formation and this mode of interaction with several of its partners appear to exert constraints on its evolution. Although, in protein-protein interaction the interacting surfaces co-evolve (47), any change on IGF-1 requires concomitant change on all the six binding proteins and on IGF-1R, which is difficult to expect as these changes are random in nature. In fact, almost all the residues of IGF-1 are highly conserved throughout chordates. In agreement with the above analysis, it was interesting to note that residues of IGF-1 (from the A and B chains) that are exposed in the complex are those that are not conserved (Figure 2 and Figure 4A).

**Figure 4:**
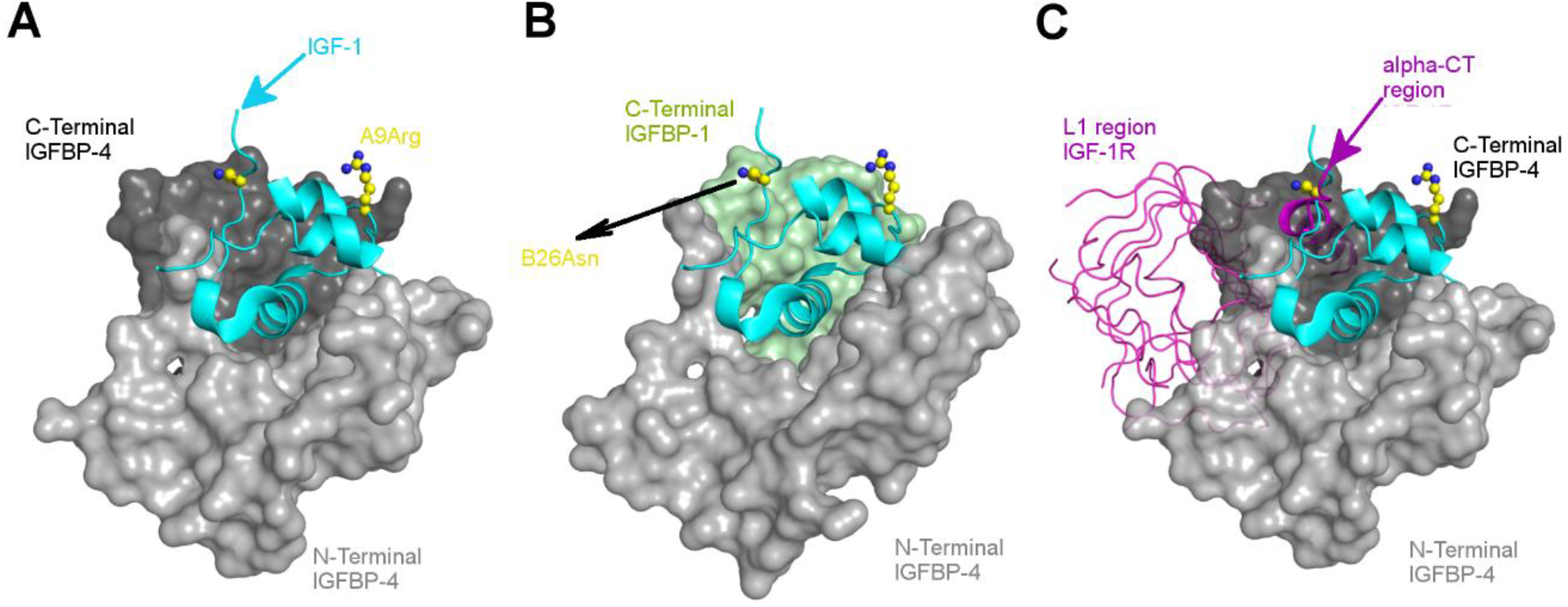
Interaction of IGF-1 with IGFBPs and IGF-1R: **A** Co-complex structure of IGF-1 with IGFBP-4 (PDB entry: 2DSR, N-terminal domain: light grey, surface representation and C-terminal domain: dark grey, surface representation). **B**: Co-complex structure of IGF-1 with IGFBP-4 and IGFBP-1 (PDB entry: 2DSQ, N-terminal domain IGFBP-4: light grey, surface and C-terminal domain of IGFBP-1: light green, surface). **C**: Superimposed structure of co-complex of IGF-1 and its receptor (5U8Q; L1 region-magenta, ribbon; αCT region-magenta cartoon) with co-complex structure of IGF-1 with IGFBP-4 (N-terminal; light grey, surface). IGF-1 is displayed in cartoon representation (cyan) across all the three panels. Note that the mode of interaction of IGF-1 with IGFBPs is very similar, where IGF-1 is bound between N and C-terminal domains of IGFBPs burying itself in a deep cleft.

Superposition of IGF-1 from the IGF-1:IGF-1R complex on to IGF-1 of IGF-1:IGFBP4 complex or IGF-1:IGFBP1/4 complex revealed that both IGFBPs and IGF-1R compete for largely a common surface on IGF-1. Further, the extent of interaction surface involved in the IGF-1:IGFBP interaction is more than that observed in the case of IGF-1:IGF-IR complex (Figure 4C). From the above analysis of the co-complex structures, it was clear that IGFBPs inhibit the interaction of IGF-1 with its receptor and hence regulate their function. Further it was observed that the α-CT region occupies some additional area other than that occupied by IGFBPs.

All IGFBPs have the capability to bind to both IGF-1 and IGF-2 and IGFBP-1 to 5 are said to have no preference between IGF-1 and IGF-2, whereas, IGFBP-6 is supposed to have higher affinity to IGF-2 than IGF-1 (48). IGF-2 is known to bind to IGF-2R which is also known as cation-independent mannose-6-phosphate receptor, with distinct sequence and domain organizations when compared to IGF-1R and IR (49). Unfortunately, there are no experimentally determined structures of IGF-2 with any of the IGFBPs to draw inferences from. However, considering the similarity of IGF-1 and IGF-2 structures, it can be envisaged that the mode of interaction between IGFBPs and IGF-2 might be comparable to that of IGF-1, and hence similar structural constraints as observed in the case of IGF-1 might be applicable to IGF-2. Thus, it appears that IGF’s resistance to sequence diversity appears to be driven by binding partner(s) imposed constraints.

The other interesting aspect is that the IGF-1 has the shortest C-peptide (12 residue) among ISP members, which is known to interact with its receptor (50). In line with this observation, C1Gly, C2Tyr, C3Gly, C7Arg and C8Arg are highly conserved (Figure S4). From the analysis of co-complex structures (PDB entry: 5U8Q) it was clearly evident that the basic residues (C7Arg and C8Arg) are important in IGF-1-IGF-1R interaction where they interact with acidic loop in the CR region, which is supported by experimental evidences (50). Further discussions can be found in the supplementary information.

### Low sequence conservation in relaxins and redefining their receptor binding motif (Relaxin 1, 2 and 3)

As observed in the phylogenetic tree (Figure 1), relaxins are distantly related to insulin and IGFs, and have very low conservation of sequence among themselves (Figure 5). Relaxins are known to perform different functions and interact with distinct GPCRs, which probably explain their sequence divergence. Further, only relaxin-2 is known to form dimers and there are no reports on dimerization of relaxin-1 or relaxin-3 (24, 27). Moreover, their function is not regulated by carrier proteins as observed in IGFs and hence their evolution is not probably regulated by partner imposed evolutionary restraints as observed in IGFs. This places less demand on the conservation of these surface exposed residues. Probably, all the reasons mentioned above contribute to the observed variation in relaxin sequences. Conversely, it is tempting to propose that the observed variation in the sequences of relaxin family hints at the possibility that they might not interact with many binding partners, unlike seen in the case of IGFs.

**Figure 5:**
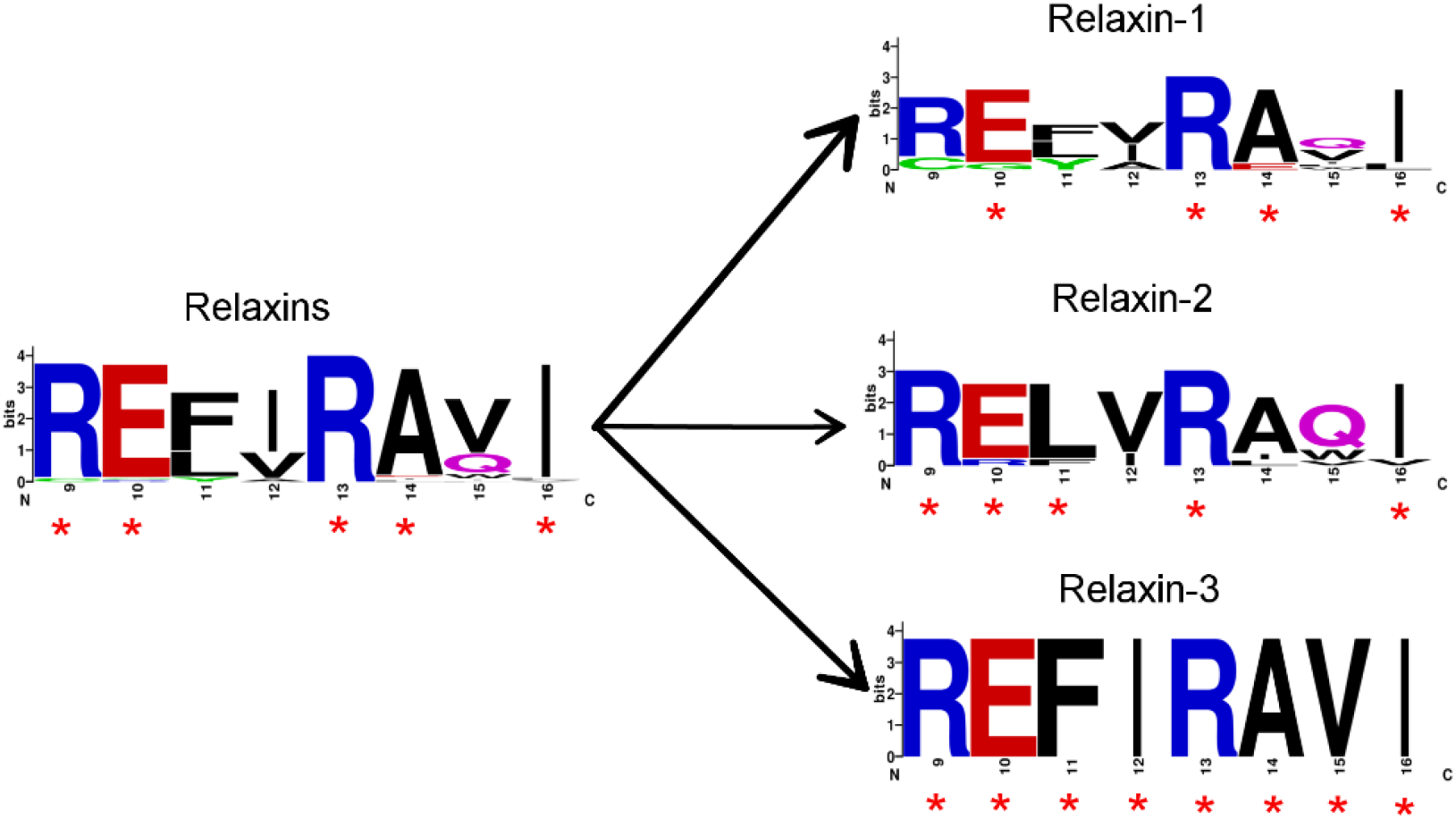
Receptor binding motif of relaxins. On the left WebLogo of receptor binding motif of relaxins (1, 2 and 3) and on the right the receptor binding motif of relaxin-1, 2 and 3 separately depicting the conservation of residues. The red asterisks represent the highly conserved residues (above 90%).

With all these sequence variations among relaxins, a conserved motif (RXXXRXXI/V) located on the B chain is shown to be important for receptor binding (51). When the receptor binding motif is analyzed separately for relaxin-1, 2 and 3, we noticed complete conservation in the receptor binding motif of relaxin-3 (**REFIRAVI**) which is not observed in relaxin-1 and 2 (Figure 5). As relaxin-3 is functionally divergent to relaxin-1 and 2, the high conservation of its receptor binding motif indicates a strict requirement for its receptor specific binding.

Surprisingly, we observed that the residues succeeding the conserved arginines (B9Arg and B13Arg) of this motif are also highly conserved among all the three relaxin paralogues. This additional conservation of B10Glu (95.9%) and B14Ala (93.9 %) in the relaxin family (relaxin-1, 2 and 3) may redefine the receptor binding motif as **RE**XX**RA**X**I/V**, instead of **R**XXX**R**XX**I/V**. Two other residues, A13Lys and B24Trp were also found to be conserved. While, B24Trp (89.1%) is known to be involved in receptor binding interaction (52), the role of highly conserved A13Lys (91.8% and 8.2% of Arg) was not mentioned before. Structural analysis reveals that A13Lys and B10Glu are in close proximity to the other two positively charged residues (B9Arg and B13Arg) of the receptor binding motif. Hence, A13Lys might also have a role in receptor binding interactions as it forms a largely positively charged patch along with the conserved residues B9Arg and B13Arg. Thus, we propose that A13Lys may also be a key residue involved in receptor binding interactions in relaxins along with B24Trp and the receptor binding motif.

### Conclusion

Our analysis of sequence conservation/variation, which stems from the phylogenetic analysis and the information from functional and structural studies, provide unprecedented insights. For example, rationale for unexpectedly high evolutionary conservation of IGFs presented here, takes into account, the function of IGFs, their regulation by IGFBPs and structural information that decipher the mode of interaction of IGFs with IGFBPs and IGF-1R. Similarly, novel findings such as (a) non-canonical C-peptide cleavage site in killifish insulin sequences (b) redefined signature motif of relaxin family, all of which are a result of collective analysis of phylogeny, sequence together with their structural and functional information. Further, in the absence of critical information regarding bombyxin receptor or its interaction with bombyxin, the sequence conservation alone suggests, unlike other insulin family members, bombyxin probably interacts with its receptor majorly utilizing its A chain. We were surprised to learn, how such an interlinked analysis that exploits information from multiple disciplines provide many profound insights on this well studied superfamily.

Insulin family proteins are known to evolve from a single ancestral gene and these Lilliputians of the protein world are the regulators of many physiological processes in higher organisms, including sugar metabolism, growth and reproduction. A recent discovery (53) reveals that the fairly new member of this family, relaxin-3, is a neuropeptide and a potential target for the treatment of various neuropsychiatric disorders (54). It is intriguing to note that nature has optimized these sequences to perform optimal and intended function over millions of years, encompassing species of diverse nature and habitat. It appears that it would be apt to borrow this wisdom from nature rather than engaging in high throughput experiments for the modification of these molecules. For example, Insulin analogue “insulin lispro” where the natural swapping of residues at the end of B-chain of IGF-1(-B28-B29-) with respect to insulin, is borrowed and incorporated on to insulin to create therapeutically successful fast acting analogue of insulin. Similar mutations can be strategized by learning from other members of the insulin superfamily for production of better insulin analogues. Intuitive designing of proteins by introducing characteristic mutations inspired from other members of this protein family might lead to potential therapeutics like dual-hormone Insulaxin (28). Conversely, this comprehensive study also points to how one can learn from nature to slightly manipulate the sequence of the protein without loss of primary function probably devoid of any cross reactivity.

## Materials and methods

### Sequence identification and analysis

Insulin family protein sequences were obtained from UniProt database (55) by using InterPro ID: IPR022352 and the clusters of each ISP members were obtained by giving 50% sequence identity cut off using UniRef database. The clusters thus obtained which consisted of at least 10 sequences were considered for the study. Some of the sequences were manually removed as they either contained incomplete sequence or were annotated as uncharacterized protein. CD-HIT server (56) was used to remove redundant dataset having 100% sequence identity by giving a higher identity cut-off of 99%, which gave 463 sequences (consisting B, C, D and E domains; for conventions of naming these domains see Figure S1) were obtained and utilized to perform multiple sequence alignment (MSA) using Clustal Omega (57). Out of the 463 sequences, insulin clade had 175 sequences, IGF-1-98, IGF-2-109, Bombyxin-32 and relaxin-49 sequences. Signal peptide was manually removed from all the sequences. MSA visualization, generation of consensus sequence and extraction of A, B and C chains were performed using Geneious (58). The images for multiple sequence alignment was generated using ENDscript server (59). The average sequence identity and most distant pair was determined using ALISTAT server (http://caps.ncbs.res.in/iws/alistat_ali.html). Residue numbering is in accordance with the numbering of master MSA sequence of processed A chain and B chains. Residues are considered to be conserved if they exhibit a conservation of >85% in the data set.

### Phylogenetic analysis

The evolutionary history was inferred using the Minimum Evolution (ME) method (60). The optimal tree with the sum of branch length = 20.99532940 is shown. Rigorous bootstrapping (500 replicates) was performed. The tree is drawn to scale, with branch lengths in the same units as those of the evolutionary distances used to infer the phylogenetic tree. The evolutionary distances were computed using the Poisson correction method (61) and are in the units of the number of amino acid substitutions per site. The ME tree was searched using the Close-Neighbor-Interchange (CNI) algorithm (62) at a search level of 1. The Neighbor-joining algorithm (63) was used to generate the initial tree. All positions with less than 95% site coverage were eliminated. Evolutionary analyses were conducted in MEGA6 (64). Phylogenetic tree was visualized by FigTree (http://tree.bio.ed.ac.uk/software/figtree/). Further, MSA for individual clades representing the families and few major paralogs as in the case of IGFs were obtained.

### Structural analysis

The protein structures of interest obtained from the world wide Protein Data Bank (wwPDB) were visualized using Pymol (65) and COOT (66). The PDB entry of proteins which were used for the study are as follows; insulin: 1ZNJ, 1EVR, 4F8F and 5BTS, IGF-1: 1GZR, IGF-2: 3KR3, relaxin-3: 2FHW, relaxin-2: 2MV1 and 6RLX and Bombyxin: 1BOM, IGF-1 binding proteins in complex with IGF-1: 2DSR, 2DSQ, Insulin receptor in complex with insulin: 4OGA, IGF-1 receptor in complex with IGF-1: 5U8Q.

PDBe PISA server was used for the analysis to decipher the interacting residues in protein complexes and in the insulin oligomer (67). This was accomplished by analyzing the buried surface area between the protein complexes and oligomers. The buried surface area of IGF-1 upon interaction with the IGFBPs was calculated using CCP4’s AreaImol (68).

## Author contributions

S.S, S.V.K, D.D, A.K, U.A.R. together conceptualized the problem. S.S. and S.V.K performed the analysis. S.S, S.V.K, A.K, U.A.R together evaluated the analysis and wrote the manuscript.

## Acknowledgments

SS would like to thank Admar mutt education foundation (AMEF) and Kongovi foundations for fellowship.

