## Supplementary material for "Phylogenetic, sequence and structural analysis of Insulin superfamily proteins reveals an indelible link between evolution and structure-function relationship"

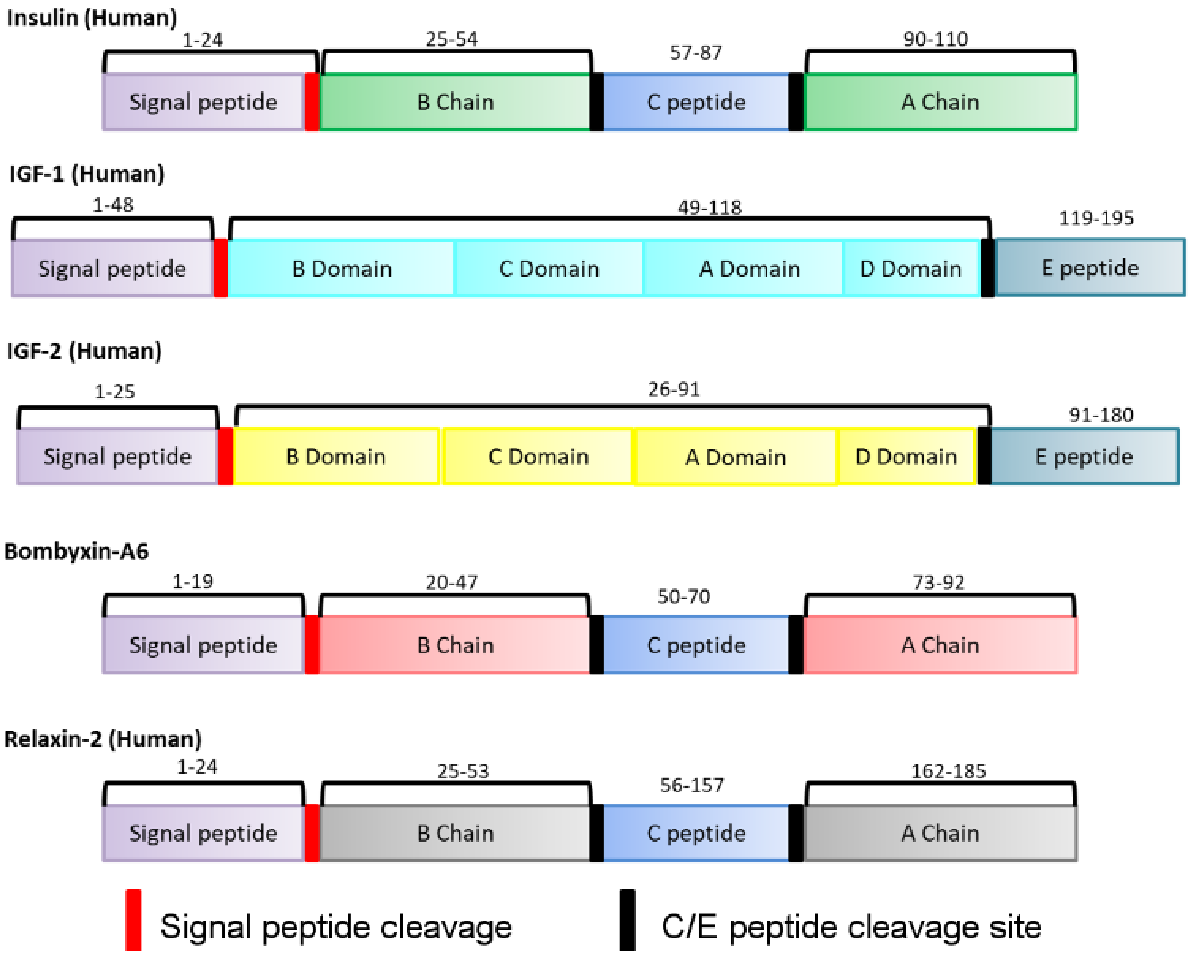


**Figure S1: Processing sites of insulin superfamily proteins.** Reference sequences have been used for the representation. The domain architecture and the signal peptide (red bar) and C/E peptide cleavage sites (black bar) have been represented.

**Table S1: Sequence identity of seven insulin superfamily proteins**

| | Insulin (P01308) | IGF-1 (P05019) | IGF-2 (P01344) | Bombyxin (P26729) | Relaxin-1 (P04808) | Relaxin-2 (P04090) | Relaxin-3 (Q8WXF3) |
|---|---|---|---|---|---|---|---|
| **Insulin** | 100 | 29.8 | 28.7 | 29.5 | 16.5 | 14.2 | 19.7 |
| **IGF-1** | 29.8 | 100.0 | 63.0 | 29.8 | 23.8 | 24.2 | 15.0 |
| **IGF-2** | 28.7 | 63.0 | 100.0 | 31.7 | 12.9 | 11.3 | 16.9 |
| **Bombyxin** | 29.5 | 29.8 | 31.7 | 100.0 | 14.1 | 13.7 | 22.9 |
| **Relaxin-1** | 16.5 | 23.8 | 12.9 | 14.1 | 100.0 | 80.4 | 24.5 |
| **Relaxin-2** | 14.2 | 24.2 | 11.3 | 13.7 | 80.4 | 100.0 | 23.0 |
| **Relaxin-3** | 19.7 | 14.3 | 16.9 | 22.9 | 24.5 | 23.0 | 100.0 |

The percentage sequence identity observed between the different members of the family without their signal peptide are tabulated above (Uniprot IDs are given in parenthesis). Note that except for the Bombyxin (*Bombyx mori*) all other sequences are from human.

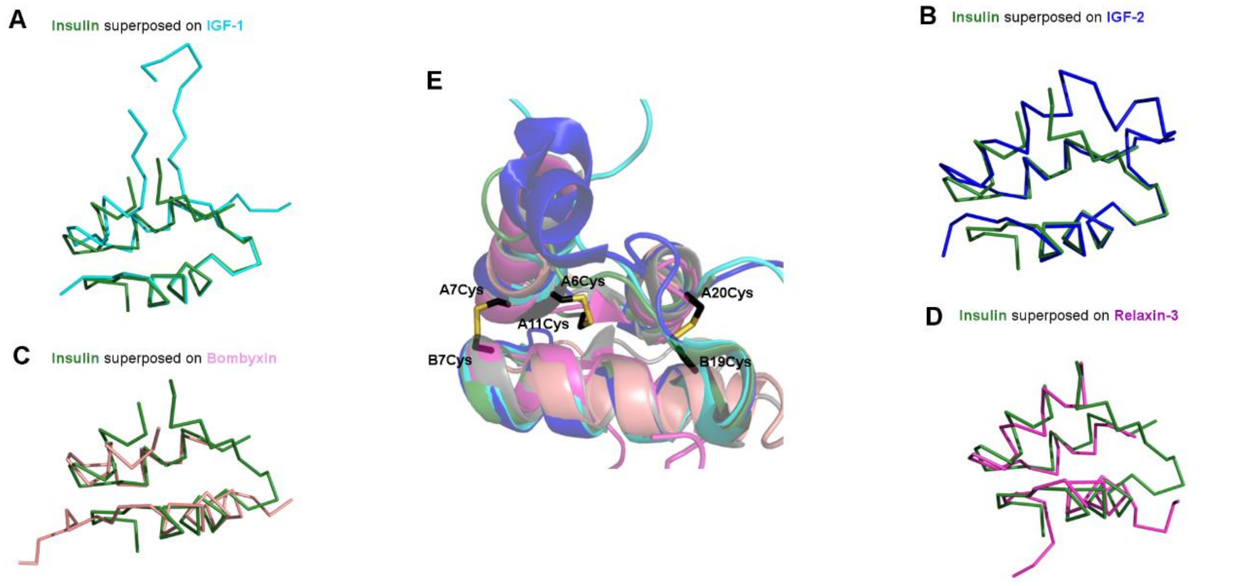


**Figure S2: The structural architecture of ISPs** consisting of three uniquely arranged helices forming the "insulin fold" have been represented. **A**: Superposed insulin (Green) and IGF-1 (Cyan) in ribbon presentation. **B**: Superposed insulin (Green) and IGF-2 (Blue) in ribbon presentation. **C**: Superposed insulin (Green) and Bombyxin (Salmon) in ribbon presentation. **D**: Superposed insulin (Green) and Relaxin-3 (Pink) in ribbon presentation. **E**: The superimposition of insulin and the reference proteins have been depicted in the middle where the canonical conserved cysteines are represented in sticks (black and yellow). The insulin fold is maintained in all the proteins except for few N/C - terminal truncations or extensions.

**Table S2: Hexamer potential of Insulin killifish sequences** based on buried surface area analysis conducted for insulin hexamer (human)

| Residue comparison between human insulin and killifish insulin for hexamer potential | | | | | | | | | | | |
|---|---|---|---|---|---|---|---|---|---|---|---|
| Human insulin | | | | Turquoise killifish (A0A1A8ACH4) | | | | Beira killifisht(A0A1A8ISB4) | | | |
| A chain | | B chain | | A chain | | B chain | | A chain | | B chain | |
| Dimer | Hexamer | Dimer | Hexamer | Dimer | Hexamer | Dimer | Hexamer | Dimer | Hexamer | Dimer | Hexamer |
| E4 (10%) | V3 (30%) | F1 (30%) | F1-10% | E4-**P** | V3 -**A**-L | F1-**A**-T | F1-**A**-T | E4-**P** | V3 -**A**-L | F1-**A**-T | F1-**A**-T |
| T8 (10%) | T8 (10%) | Q4 (10%) | V2-50% | T8-**A**-D | T8-**A**-D | Q4-**P** | V2-**A**-S | T8-**A**-D | T8-**A**-D | Q4-**P** | V2-**A**-S |
| I10 (10%) | S9 (10%) | G8 (30%) | H5-10% | I10-**A**-P | S9 -**A**-R | G8 -**P** | H5-**A**-K | I10-**A**-P | S9 -**A**-R | G8 -**P** | H5-**A**-Q |
| S12 (40%) | I10 (10%) | S9 (70%) | G8-40% | S12-**A**-T | I10 -**A**-P | S9 -**A**-P | G8-**P** | S12-**A**-T | I10 -**A**-P | S9 -**A**-P | G8-**P** |
| L13 (20%) | N21 (10%) | V12 (100%) | H10-50% | L13-**A**-I | N21 -**A**-F | V12 -**A**-T | H10-**P** | L13-**A**-I | N21 -**A**-F | V12 -**A**-T | H10-**P** |
| Y14 (30%) | | E13 (60%) | L17-40% | Y14-**A**-F | | E13 -**A**-D | L17-**P** | Y14-**A**-F | | E13 -**A**-D | L17-**P** |
| Q15 (20%) | | Y16 (70%) | V18-20% | Q15--**A**-D | | Y16 -**P** | V18-**P** | Q15--**A**-D | | Y16 -**P** | V18-**P** |
| E17 (10%) | | L17 (10%) | R22-10% | E17 -**A**-Q | | L17 -**P** | R22-**P** | E17 -**A**-Q | | L17 -**P** | R22-**P** |
| | | G20 (20%) | K29-10% | | | G20 -**P** | K29-**A**-R | | | G20 -**P** | K29-**A**-R |
| | | E21 (30%) | | | | E21 -**P** | | | | E21 -**P** | |
| | | G23 (40%) | | | | G23 -**P** | | | | G23 -**P** | |
| | | F24 (60%) | | | | F24 -**P** | | | | F24 -**P** | |
| | | Y26 (60%) | | | | Y26 -**A**-F | | | | Y26 -**A**-F | |
| | | K29 (10%) | | | | K29-**A**-R | | | | K29-**A**-R | |

Buried surface area (BSA) analysis was carried out in PDBe PISA (1) on human insulin using the structures 5BTS, 4F8F and 1EVR. The buried surface is represented inside the parenthesis and the residues have been represented in the single letter code. The residues which are present in the sequence are displayed as P and the residues which are absent are displayed as A and the residue to which it is mutated to is represented in one letter code.

**Table S3: Receptor binding residues in Killifish sequences**.

| **Site 1** | | **Site 2** | |
|---|---|---|---|
| Gly A1 | Present | Thr A8 | Mutated to Asp |
| Ile A2 | Present | Ile A10 | Mutated to Phe |
| Val A3 | Mutated to Leu | Ser A12 | Mutated to Thr |
| Glu A4 | Present | Leu A13 | Mutated to Ile |
| Tyr A19 | Mutated to Phe | Glu A17 | Mutated to Glu |
| Asn A21 | Mutated to Phe | His B10 | Present |
| Gly B8 | Present | Glu B13 | Mutated to Asp |
| Ser B9 | Mutated to Pro | Leu B17 | Present |
| Leu B11 | Present | | |
| Val B12 | Mutated to Thr | | |
| Tyr B16 | Present | | |
| Phe B24 | Present | | |
| Phe B25 | Mutated to Leu | | |
| Tyr B26 | Mutated to Phe | | |

The residues which take part in receptor binding in insulin (both site 1 and site 2) (2) have been tabulated and it was checked if the killifish insulin sequences had all the residues required for the receptor binding. The residues which are present  and the residues which are mutated are mentioned.

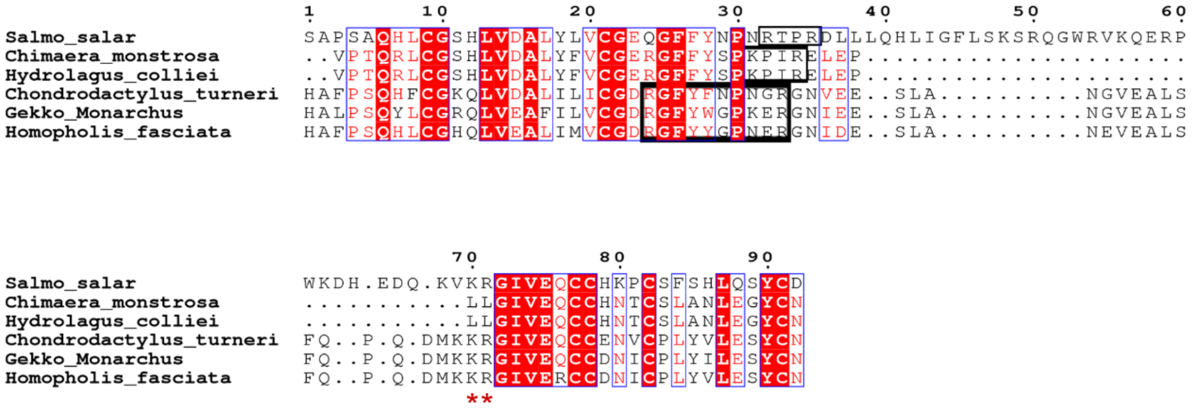


**Figure S3: Non-canonical cleavage sites** (Type 3 precursor cleavage motif) in other insulin sequences. The motif is highlighted with a black outline for each sequence.

Type 3 precursor cleavage site was checked in the entire insulin clade containing insulin sequences other than above mentioned killifish sequences. Among 175 insulin sequences studied only 6 of them had a type 3 precursor cleavage motif. Interestingly, 3 of these insulin sequences belonged to fish species and the other three belonged to gecko species (Figure S3). The sequences with the type 3 precursor cleavage site are S*almo salar*- Atlantic salmon (Uniprot ID: A0A1S3SVH7), *Chimaera monstrosa* – Rabbit fish (Uniprot ID: P68991), *Hydrolagus colliei* - *Spotted ratfish* (Uniprot ID: P68992), *Chondrodactylus turneri* – Turner's thick-toed gecko (Uniprot ID:A0A0M4TUN1), *Gekko monarchus* - spotted house gecko (Uniprot ID: A0A0M4UV57), *Homopholis fasciata* - banded velvet gecko or striped velvet gecko (Uniprot ID: A0A0M4UT51). The red asterisks represent the N-terminal cleavage site of A chain and we observed a non-canonical cleavage site again where in two sequences (*Chimaera monstrosa* and *Hydrolagus colliei*) di-hydrophobic residues (leucine) are present. The rationale behind this change in the cleavage site is not known. Hence, this rare type 3 precursor cleavage motif was found to be predominantly present in fishes than any other species.

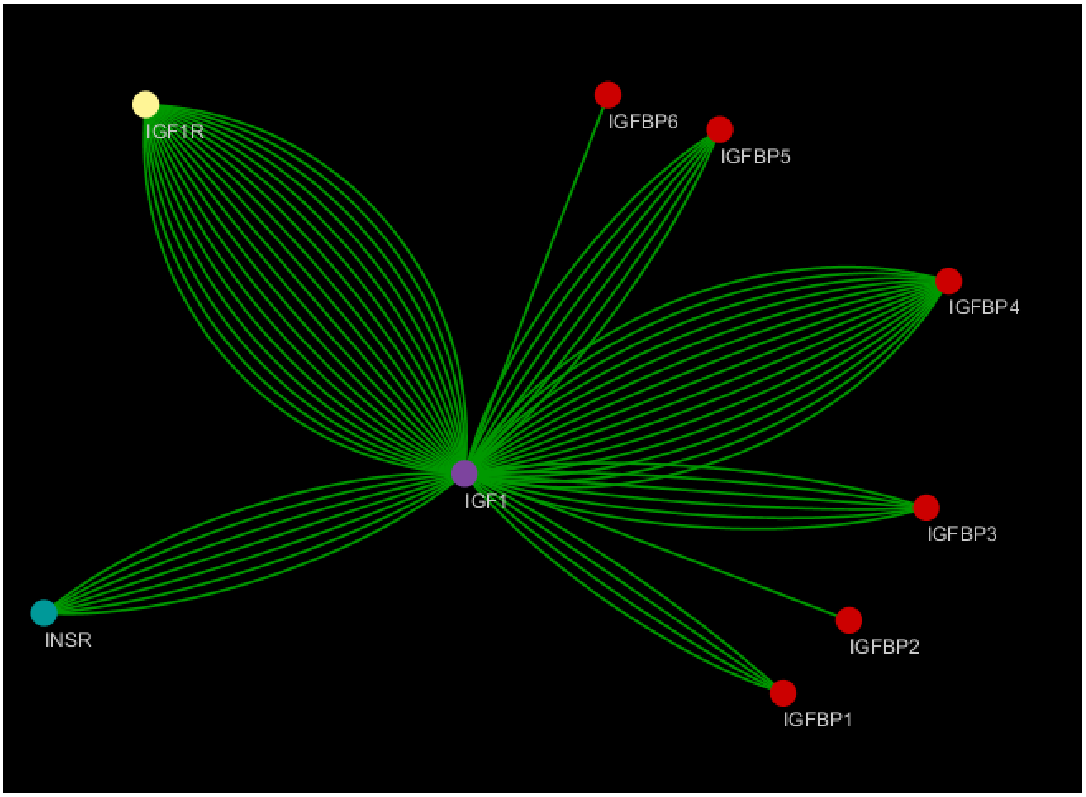


**Figure S4: A simple interactome representing the interactions of IGF-1 with its partners**
All the interactions with experimental studies have been represented by green lines where IGF-1 (purple circle) is the central node which interacts with other proteins. The red circles represent the IGFBPs (1-6), the blue circle represents insulin receptor, and the yellow circle represents the IGF-1 receptor. Cystoscape was used to construct the protein interactome of IGF-1 with its binding partners and the database used to ascertain the experiments performed to establish the interactions was iRefIndex (3).

**Conserved arginines in the C peptide of IGF-1 provide additional affinity in IGF-1:IGF-1R complex**

The C-peptide of IGFs, in particular IGF-1, is distinct from other family members. Here the C-peptide is the shortest among ISPs (12 residues in IGF-1 and 16 residues in IGF-2), is not cleaved post-translationally, and has relatively high sequence conservation compared to other ISP family members. Our sequence analysis indicates that residues C1Gly, C2Tyr, C3Gly, C7Arg and C8Arg are almost completely conserved in IGF-1 (Figure S5C). In general, it can be expected that such a high conservation during the evolution has some functional role. In line with this observation, in human IGF-1, the conserved arginine residues C7Arg, C8Arg, has been known to make vital salt-bridge interactions with the acidic patch of the CR region (289Glu, 292Asp and 294Glu on human IGF-1R, Uniprot ID: P08069), which is determined biochemically (4).

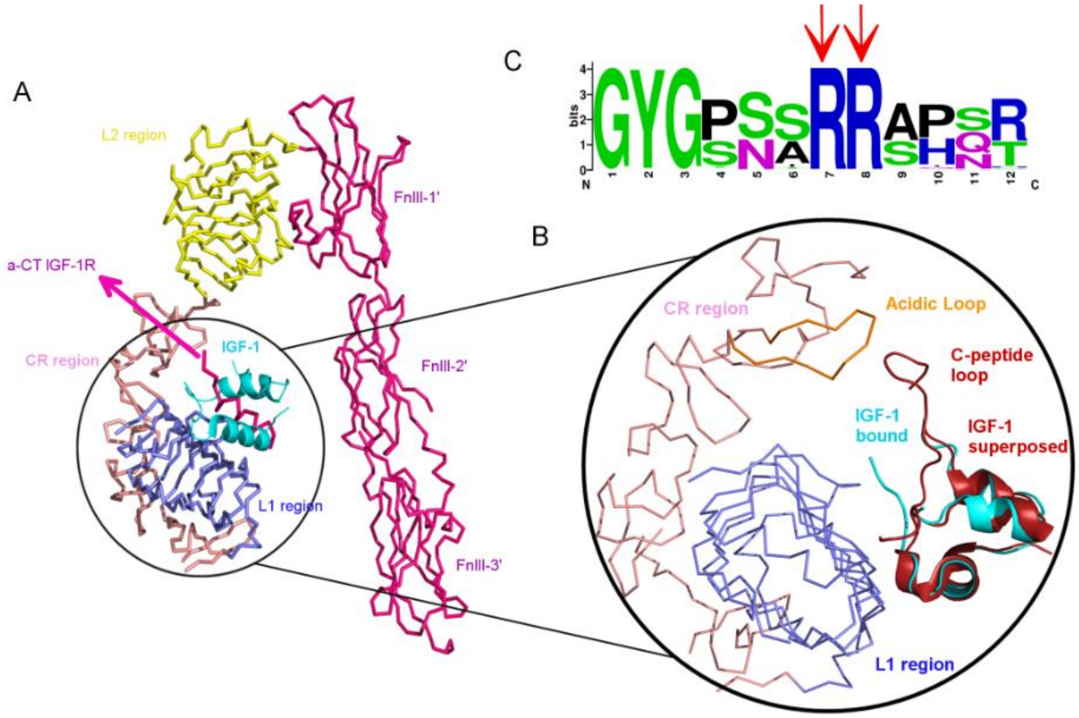


**Figure S5: Interaction of IGF-1 with its receptor (IGF-1R). A:** Co-complex structure of IGF-1 with its receptor IGF-1R (PDB ID: 5U8Q). Different domains of the IGF-1R are colored differently and labeled. Note that IGF-1 interacts with Leucine rich region L1 (slate-blue, ribbon) and the C-terminal helix called α-CT region (pink, ribbon) **B:** Superimposed structures of IGF-1:IGF-1R complex (5U8Q) with IGF-1 (1GZR) with C-peptide. It is clear that the C-peptide is in close proximity to CR-region containing acidic patch. **C:** Sequence logo of IGF-1 C peptide residues highlighting the conservation of residues -C1Gly-C2Tyr-C3Gly- and two arrows pointing the conserved -C7Arg-C8Arg- residues.

Ectodomains of insulin receptor (IR) and insulin-like growth factor-1 receptor (IGF-1R) share similar structure and domain architecture, with a leucine rich region 1 (L1), a cysteine rich region (CR), a Leucine rich region 2 (L2) and an insert domain followed by C-terminal helix called α-CT region (Figure S5a). Based on the biochemical and biophysical experiments it has been shown that both insulin and IGF-1 have major interactions with the L1 region and α-CT region of the respective receptors. Recent structure of IGF-1 in complex with IGF-1R unequivocally supports the above observation (5). However, since this structure was determined by soaking the IGF-1 with native IGF-1R crystals, the electron density for the C-peptide region is not clear. Superposition of IGF-1 structure containing C-peptide loop (PDB-entry:1GZR) on to IGF-1 bound to IGF-1R (PDB-entry: 5U8Q) suggest that the conserved arginine residues on the C-peptide of IGF-1 are in close proximity to acidic patch in the CR region of IGF-1R. The observed evolutionary conservation of consecutive arginine residues on the C-peptide of IGF-1 in our analysis corroborates well with the proposition that these arginines make key interactions in the IGF-1R:IGF complex (6). This is further supported by a report on IGF-1 mutant with the replacement of entire C-peptide region of IGF-1 with four consecutive glycine residues resulting in a 30 fold loss of affinity to its receptor IGF-1R (6). The absence of these salt bridge interaction in the case of IGF-1:IR complex probably contributes to the observed low affinity cross-reactivity.
